# TNFα increases Tyrosine Hydroxylase expression in human monocytes

**DOI:** 10.1101/2021.04.13.439627

**Authors:** Adithya Gopinath, Martin Badova, Madison Francisa, Gerry Shaw, Anthony Collins, Douglas R. Miller, Carissa A. Hansen, Phillip Mackie, Malú Gámez Tansey, Abeer Dagra, Irina Madorsky, Adolfo Ramirez-Zamora, Michael S. Okun, Wolfgang J. Streit, Habibeh Khoshbouei

## Abstract

Most, if not all, peripheral immune cells in humans and animals express tyrosine hydroxylase (TH), the rate limiting enzyme in catecholamine synthesis. Since TH is typically studied in the context of brain catecholamine signaling, little is known about changes in TH production and function in peripheral immune cells. This knowledge gap is due, in part, to the lack of an adequately sensitive assay to measure TH in immune cells expressing lower TH levels compared to other TH expressing cells. Here, we report the development of a highly sensitive and reproducible Bio-ELISA to quantify picogram levels of TH in multiple model systems. We have applied this assay to monocytes isolated from blood of persons with Parkinson’s disease (PD) and to age-matched, healthy controls. Our study unexpectedly revealed that PD patients’ monocytes express significantly higher levels of TH protein in peripheral monocytes relative to healthy controls. Tumor necrosis factor (TNFδ), a pro-inflammatory cytokine, has also been shown to be increased in the brains and peripheral circulation in human PD, as well as in animal models of PD. Therefore, we investigated a possible connection between higher levels of TH protein and the known increase in circulating TNFδ in PD. Monocytes isolated from healthy donors were treated with TNFδ or with TNFδ in the presence of an inhibitor. Tissue plasminogen activator (TPA) was used as a positive control. We observed that TNFδ stimulation increased both the number of TH+ monocytes and the quantity of TH per monocyte, without increasing the total numbers of monocytes. These results revealed that TNFδ could potentially modify monocytic TH production and serve a regulatory role in peripheral immune function. The development and application of a highly sensitive assay to quantify TH in both human and animal cells will provide a novel tool for further investigating possible PD immune regulatory pathways between brain and periphery.

## Introduction

Human and animal studies have shown that most if not all immune cells possess components necessary to release, uptake, synthesize, and respond to catecholamines including dopamine and norepinephrine (NOR). These components activate signaling cascades that change the phenotype and function of cells in both healthy and in disease conditions. Immune cells may thus both come in contact with physiological levels of catecholamines derived from peripheral tissues and also serve as a source for catecholamines. Tyrosine hydroxylase (TH) catalyzes the conversion of tyrosine to 3,4-dihydroxyphenylalanine (L-DOPA), which is the rate-limiting step in the synthesis of dopamine, norepinephrine (NOR) and epinephrine^1, 2^. Although primarily studied in the central nervous system^3, 4^, TH is expressed in the majority of peripheral immune cells^5–9^, and many peripheral tissues^10^, including kidney^11, 12^, heart^13^ and adrenal cortex^14–16^. Both myeloid and lymphoid lineages of human peripheral immune cells express TH^17, 18^, which is thought to regulate dopamine levels within these cells^9^. Beyond protein expression, TH activity is regulated by a variety of post-translational modifications and that can regulate TH function. For example, phosphorylation, ubiquitination, nitration and S-glutathionlyation can all affect TH activity independent of TH levels^19–26^. As the key to catecholamine production, TH activity and its relative expression are commonly studied in diseases in which catecholamine tone, synthesis and signaling are altered. These disease states include bipolar disorder, addiction, schizophrenia, attention deficit hyperactivity (ADHD) and neurodegenerative conditions including Parkinson’s disease (PD).

The lack of a robust and sensitive assay to measure low levels of TH protein has hampered the field’s ability to investigate TH protein levels in peripheral immune cells in diseases characterized by altered catecholamine tone. For example, in PD, due to its spatially restricted expression, decreases in TH levels in the basal ganglia are readily detectable^27, 28^, whereas changes in TH levels in other brain regions (i.e., amygdala, hippocampus, cortical regions) are reported in the later stages of PD^29, 30^. In contrast, very low TH levels in countless immune cells spread across the body has made it difficult to study TH protein levels in peripheral immune cells. For example, indirect TH measurements via qPCR reveal that PD patients show significantly less midbrain TH mRNA compared to healthy controls subjects (5.5 ± 1.4 in healthy controls, vs. 1.5 ± 0.9 attomole/microgram total RNA in PD)^31^. In contrast, TH mRNA is not detectable in unstimulated immune cells^32^. TH protein expression in the substantia nigra is in excess of 200ng TH per milligram protein^33^, and is decreased in patients with PD. However, to our knowledge no reports directly quantify TH protein in immune cells.

In order to investigate whether the characteristically reduced TH expression in PD central nervous system (CNS) is recapitulated in peripheral immune cells, we established a sensitive assay to quantify TH protein. We then applied the assay to analyze TH production in peripheral blood monocytes. The sensitivity of our Bio-ELISA was a thousand-fold above traditional detection methods, and when we measured TH level in peripheral monocytes from healthy controls and from PD, we observed a significant elevation of TH levels in PD monocytes versus controls. This observation was contrary to our *a priori* hypothesis. The unexpected discovery of increased TH protein in peripheral PD monocytes prompted investigation into the potential underlying mechanism. In the PD literature, there is a strong consensus that neuroinflammatory cytokines, including TNFδ and IL6 are increased in CSF and serum of PD patients and of animal models of PD^27, 34–42^. Therefore, we investigated whether *ex vivo* exposure to TNFδ or IL6 increases the number of TH+ monocytes and/or amount of TH protein per monocyte. We found that exposure to TNFδ, but not IL6 increased both the number of TH+ monocytes and the quantity of TH protein per cell.

## Results and discussion

### Bio-ELISA successfully and reproducibly detects recombinant and native TH

To test the hypothesis that similar to CNS in PD, TH expression is reduced in peripheral blood monocytes, we first established a sensitive assay to quantify TH levels in monocytes from healthy controls, as well as various reference TH expressing systems. Given the plethora of biological systems expressing TH, there is an unmet need for a sensitive and reliable assay to quantify TH levels which with broad biological implications in basic science, preclinical and clinical research. To date, measurement of TH levels in midbrain neurons has been accomplished by immunohistochemistry, and Western blot^62–65^, while TH levels in peripheral immune cells has been assayed by flow cytometry^51^. Although reliable, these methods share a common shortcoming in that they are semi-quantitative at best, and at worst only indicate the presence or absence of TH. This led us to develop a highly sensitive and fully quantitative enzyme-linked immunosorbent assay (Bio-ELISA) to measure TH protein levels.

Quantification of TH using Bio-ELISA depends on the availability of purified TH and high-quality antibodies against TH, preferably generated in two distinct host species. A panel of monoclonal and polyclonal antibodies were generated against full length recombinant human TH (Figure 1A), and quality assessment was performed by standard ELISA, Western blotting, and appropriate cell and tissue staining. These novel antibodies behaved in all respects similarly to a widely used commercial TH antibody (Figure 1B, AB152, Millipore-Sigma)^66–69^. A mouse monoclonal antibody, MCA-4H2, and a rabbit polyclonal, RPCA-TH, were selected as ELISA capture and detection antibodies, respectively.

**Figure 1.**
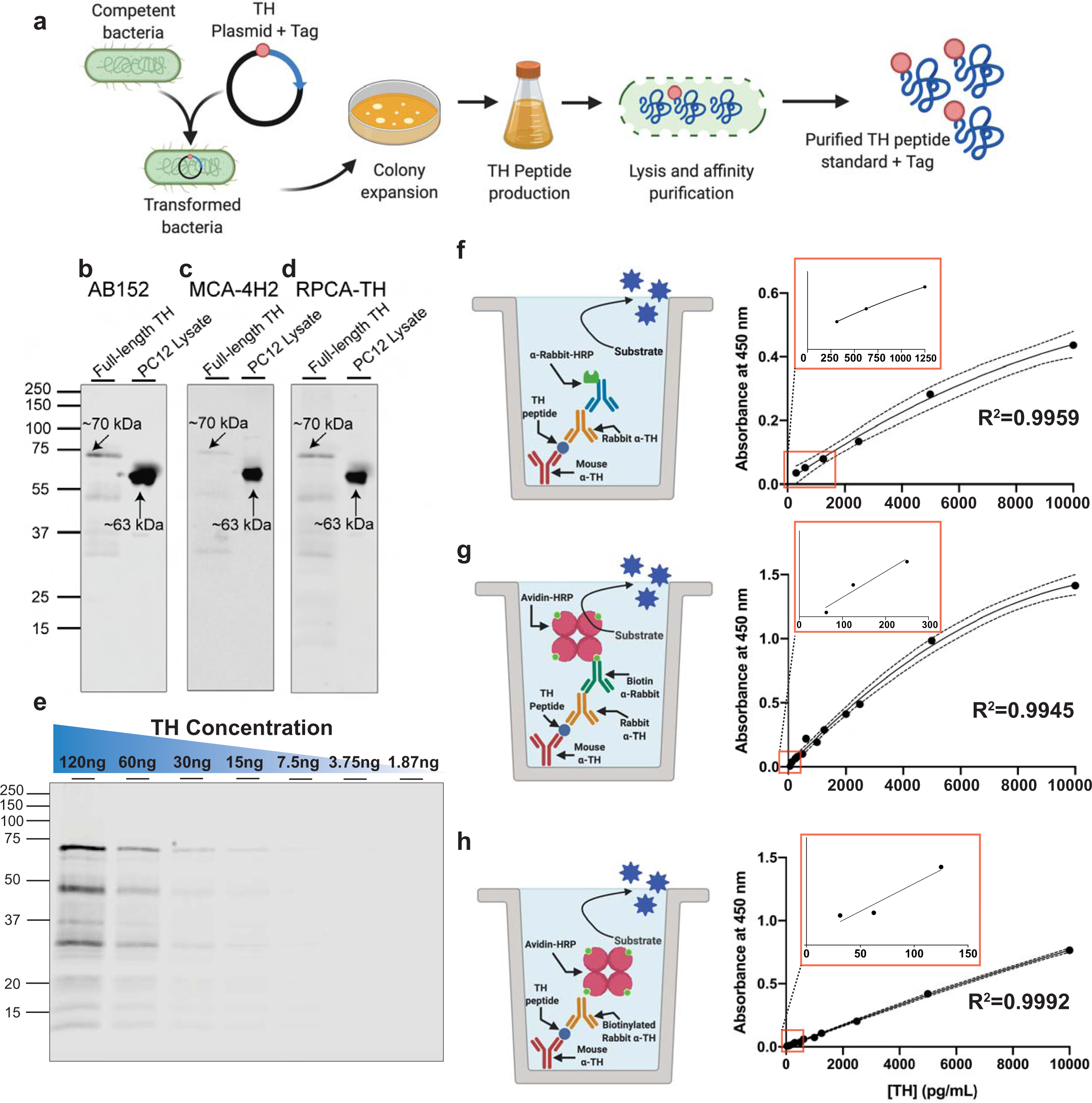
Establishing a reproducible quantitative Bio-ELISA to detect Tyrosine Hydroxylase. A-D) TH is detectable in recombinant form and in PC12 crude lysate using affinity purified rabbit polyclonal TH antibody AB152 (Sigma), and antibodies selected for this ELISA, mouse monoclonal MCA-4H2 (EnCor) and rabbit polyclonal RPCA-TH (EnCor). E) Using AB152, we probed the lower threshold for TH detection *via* serial dilution of purified recombinant TH from 6ug/mL to 0.094ug/mL followed by Western blot and near-Infrared detection, considered to be a sensitive method for protein detection on Western blot. We demonstrate IR detection is reliable to a lower threshold of ∼15ng TH. Below this limit, TH detection becomes unreliable with IR detection. F-H) In a series of stepwise experiments designed to increase ELISA sensitivity and decrease background, we achieved lower detection limits of 15pg/mL TH (H). Capture antibody and detection antibody in all three methods were MCA-4H2 (1:1,000 dilution from 1mg/mL) and RPCA-TH (1:6,000 dilution from 1mg/mL). Schematic representation of each method shown on the left with representative standard curve on the right. F) Incubation with detection antibody followed by an HRP-conjugate secondary yielded a lower detection threshold of 125pg/mL. G) Addition of a tertiary layer using anti-rabbit biotin followed by Avidin-HRP improved lower detection threshold to 62.5pg/mL but resulted in increased background. H) Use of biotinylated detection antibody (RPCA-TH-biotin, 1:6,000 dilution from 1.65mg/mL) followed by avidin-HRP yielded the lowest detection threshold of 15 pg/mL, with maximum sensitivity and minimal background. F-H) Insets (red outline) shows magnified lower standard curve to illustrate sensitivity.

Next, TH recombinant protein band identity was compared to TH expression in PC12 cells (Figure 1C). As predicted, PC12 lysate shows a single TH band at ∼63 kDa, with a corresponding band for the TH recombinant protein at ∼70 kDa. The observed difference in molecular weight between TH expressed in PC12 cells and recombinant TH protein is due to the additional 5.7kDa N-terminal His-tag. Lower molecular weight bands (at 50kDa and 35kDA, Figure 1B-E, Supplementary Figure 1) represent proteolytic cleavage products of mammalian TH when expressed in a prokaryotic system. Both MCA-4H2 and RPCA-TH reliably detect both recombinant TH and native TH in PC12 lysates (Figure 1 C, D).

Since antibody specificity is crucial for developing a novel assay, we rigorously confirmed their specificity. First, MCA-4H2 and RPCA-TH were used to stain human and murine midbrain tissue (Figure 2). MCA-4H2 (Figure 2A) and RPCA-TH (Figure 2B) both showed high specificity for TH+ dopamine neurons in both human and murine tissues with no visible background. In addition, both secondary-only and isotype control staining show minimal background (Figure 2A-B, top and second panels). Lastly, both MCA-4H2 and RPCA-TH were tested via Western blot using standard immunoblotting as well as blocking peptide/absorption controls (Figure 2C-D). Both antibodies show good specificity and minimal background. CHO cells, used as the negative control since they do not express TH, show no TH band (Figure 2C). The peptide blocking/absorption control groups (Figure 2D) also show no detectable signal, further confirming specificity.

**Figure 2.**
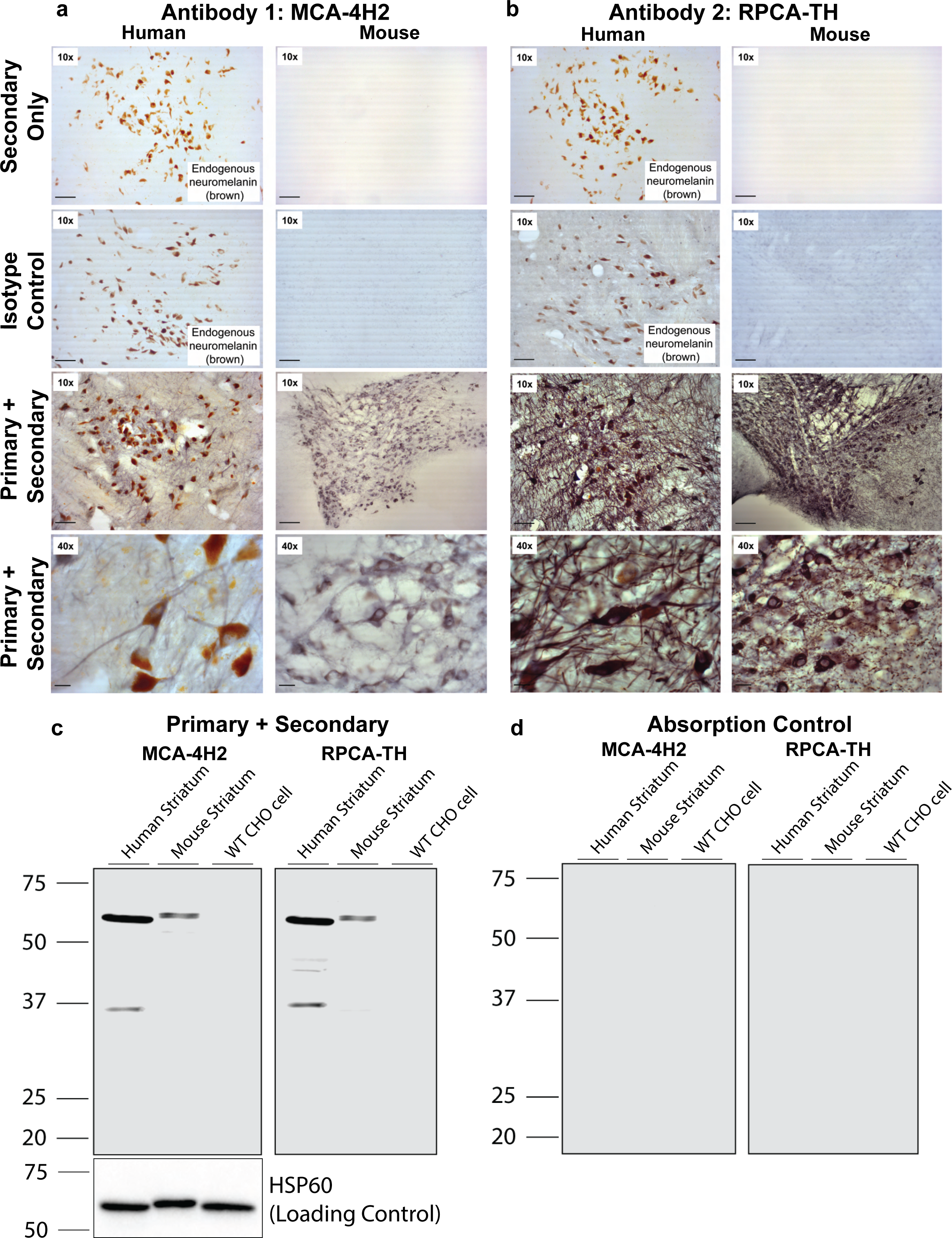
Antibodies MCA-4H2 and RPCA-TH reliably detect both native and denatured TH in mouse and human tissue. Human and murine brain sections (40um) were permeabilized, blocked and stained with primary antibodies (MCA-4H2 and RPCA-TH) followed by HRP-conjugated secondaries and detected using diaminobenzidine enhanced with nickel (NiDAB, grey-black). A) MCA-4H2 stains neuromelanin-expressing (brown) TH positive midbrain neurons and neuronal processes (grey-black) with no non-specific staining (secondary only, top panel; isotype control, second panel) in both human and murine tissues. B) RPCA-TH shows similar highly specific staining of midbrain TH positive neurons, confirming antibody specificity. A & B) Human midbrain tissues shown as secondary-only and isotype controls exhibit endogenous neuromelanin (brown), not to be confused with immunostaining. C) Western blot analyses of murine and human striatal tissues reveal similarly specific detection of TH (∼63kDa band) in both mouse and human, with minimal non-specific staining in negative control homogenate (parental CHO cell homogenate). (Left – MCA-4H2, right – RPCA-TH). D) Blocking peptide/absorption control followed by western blot detection with either RPCA-TH and MCA-4H2 confirms specificity of both antibodies for TH protein. HSP60 (loading control) is shown below and applies to C-D.

Next, we prepared 1:1 serial dilutions of TH recombinant protein in Laemmli buffer, from 6ug/mL to 0.094ug/mL, to test the limits of detection using the Licor IR imaging system for Western blot (Figure 1E) using commercially available TH antibody AB152. While effective, detection *via* Licor Odyssey using an IR fluorescent dye affords a fixed lower detection limit of ∼15ng, suggesting that IR fluorescent imaging is suitable for high expressing systems, but unsuitable for accurate quantification at low nanogram or picogram TH levels, reinforcing the need for a more sensitive, quantitative TH Bio-ELISA.

To quantify TH expression in control conditions, we first attempted a standard sandwich ELISA approach (Figure 1F), in which MCA-4H2 was used as the capture antibody, followed by incubation with recombinant TH, then RPCA-TH as detection antibody. Enzyme-based detection was accomplished by addition of HRP-conjugated secondary (goat anti-rabbit HRP, Vector, BA1000). While this reliably quantified TH, the standard version of this assay produced a lower detection threshold of 125pg/mL TH. We sought to further increase sensitivity of the assay by addition of a biotin-avidin amplification step (Avidin-HRP, Vector, A2004) (Figure 1G), which provided an improved lower threshold of 62.5pg/mL. A further refinement was the biotinylation of the rabbit detection antibody using Sulfo-NHS-LC-biotin (Thermo Scientific A39257) which improved sensitivity further by reducing background and producing a lower-threshold of detection at 15pg/mL (Figure 1H) with biotinylated antibodies, hence the Bio-ELISA designation. We found that our Bio-ELISA is around one thousand-fold more sensitive than infrared Western blot imaging (15 pg/mL vs. 15 ng/mL). Both TH antibodies are available commercially from EnCor Biotechnology Inc.

#### Antibodies MCA-4H2 and RPCA-TH reliably detect both native and denatured TH in mouse and human tissue

Aiming to develop a novel and reliable ELISA for both human and murine tissues, we next sought to confirm specificity of these antibodies on native and denatured tissues from both human and mouse brain regions rich in tyrosine hydroxylase (Figure 2). MCA-4H2 (Figure 2A) and RPCA-TH (Figure 2B) detect TH+ cell bodies and neuronal processes in both human and mouse midbrain. Minimal non-specific staining detected in secondary only and isotype controls, further confirming antibody specificity. Similarly, both MCA-4H2 and RPCA-TH detect denatured TH on Western blot (Figure 2C) following separation on SDS-PAGE, with minimal non-specific bands in the negative control (parental CHO cell homogenate). HSP60 is shown as a loading control. As an additional validation step to confirm the specificity of MCA-4H2 and RPCA-TH, primary antibodies were pre-incubated with recombinant TH protein (blocking peptide/absorption control) and show no observable signal (Figure 2D).

#### TH Bio-ELISA reliably quantifies TH in PC12 cells, human macrophages, and cultured murine dopamine neurons

Having established a reliable method with a suitably low detection threshold, we tested the TH Bio-ELISA on cell homogenates prepared from PC12 cells, HEK293 cells, cultured primary human macrophages derived from whole blood samples from healthy donors, and primary cultures of midbrain dopamine neurons prepared from PND0-PND3 mouse pups. PC12 cells are known to express high levels of TH^48^, while HEK293 serve as negative control^45, 70^. Cultured midbrain dopamine neurons are known to express TH as the rate limiting enzyme for dopamine^47^ while cultured human monocyte-derived-macrophages express TH protein and mRNA^9, 43^.

TH expression is shown as unit TH (picogram or nanogram) per mg total protein, as determined by the Lowry assay. PC12 homogenate provided a reliable positive control expressing high levels of TH (>10ng TH/mg total protein), while HEK293 homogenate showed no detectable levels of TH, in at least 6 independent replicates. As anticipated, cultured dopamine neurons from postnatal mice showed greater TH concentrations (∼700pg TH/mg total protein) than cultured human macrophages (∼300pg TH/mg total protein) (Figure 3A), suggesting the Bio-ELISA is applicable to cell and tissue samples derived from human and murine specimens, paving the way for its application in translational and preclinical studies involving measurements of TH protein. We should note that unlike cultured human monocyte-derived-macrophages, cultured dopamine neurons contain various cell types, and consist of 12-16% dopamine neurons. The remainder are GABAergic neurons and supporting cells (microglia and astroglia)^71–73^. Thus we believe that TH levels are much higher in a single dopamine neuron than in a macrophage. Visual representation of relative TH expression in PC12, HEK293, human macrophage and primary neuron homogenates are plotted on a representative standard curve (Figure 3B). Raw values [TH] in ng/mL calculated from absorbance are shown in Figure 3C, alongside each sample ID. Raw TH concentration was divided by [Protein], then multiplied by 1,000 to produce values in pg TH/mg total protein (Figure 3C).

**Figure 3.**
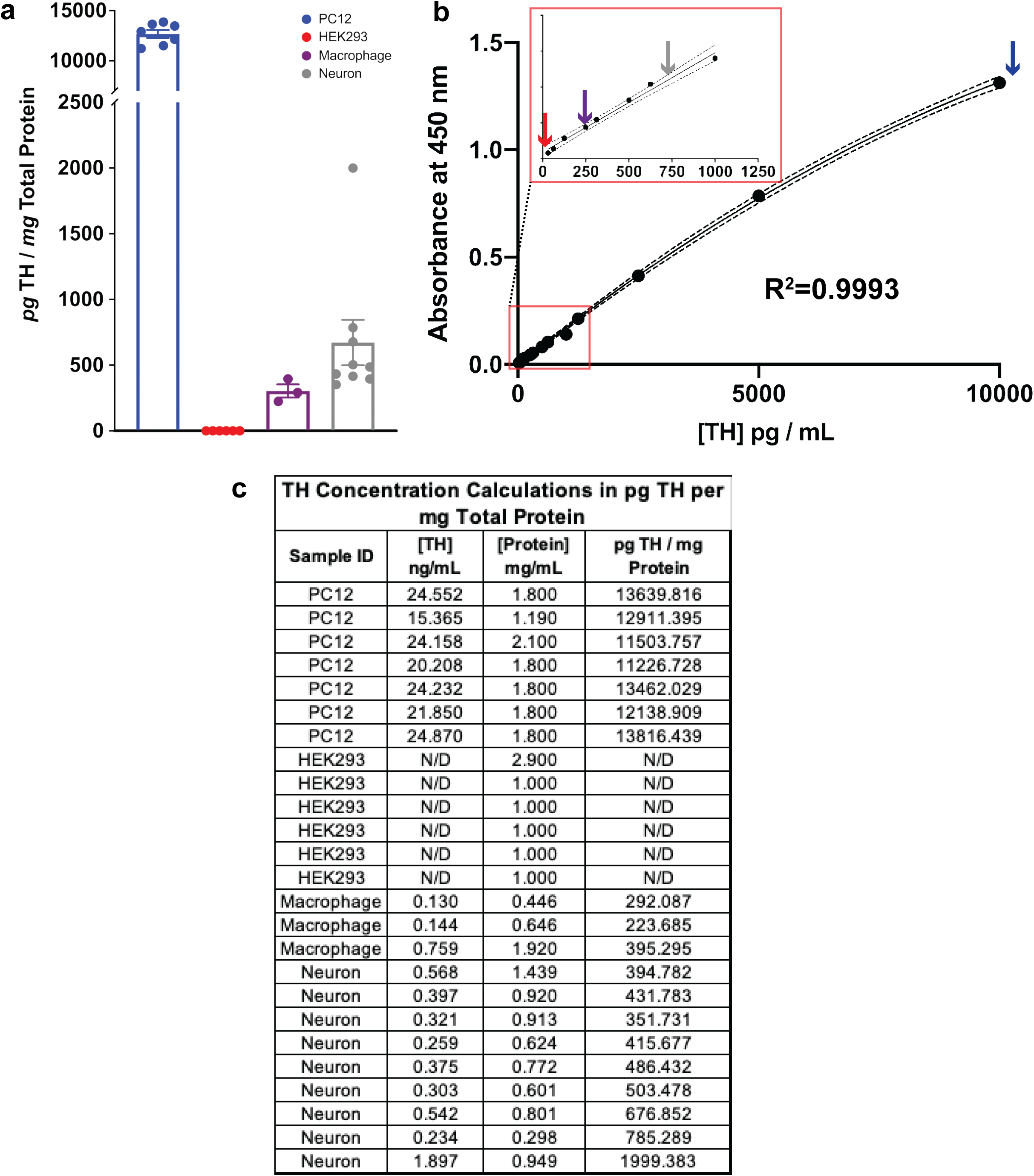
Bio-ELISA reliably quantifies TH in PC12 cells, human macrophages and cultured murine dopamine neurons. A) Using the Bio-ELISA shown in Figure 1G, we quantified TH in four relevant tissues and cultured cells: PC12 (positive control), HEK293 (negative control), cultured human macrophages and cultured primary murine dopamine neurons. PC12 cells express very high levels of TH (<10ng TH / mg total protein) relative to human macrophages (∼300 pg TH / mg total protein) and primary murine dopamine neurons (∼700 pg TH / mg total protein). B) TH values are plotted on a representative standard curve for visual comparison, with inset magnifying the lower end of the standard curve. C) Calculations are shown by which raw TH concentration in ng/mL is normalized to total protein per sample. Samples included in A, each an independent biological replicate, are shown in C. Data are shown as ±SEM.

To further confirm the specificity of these antibodies, the Bio-ELISA was tested using absorption controls (Figure 4). In multiple independent replicates, a single ELISA plate was prepared as shown in Figure 4A (Bio-ELISA, blue; absorbed MCA-4H2, orange; absorbed RPCA-TH, green), and incubated with PC12 lysate as a positive control. Following peptide blocking/absorption of either capture or detection antibody, PC12 cell lysate yields no detectable TH (orange and green arrows, Figure 4B), while the TH Bio-ELISA (blue arrow) recapitulates TH concentrations measured in PC12 cells (compare Figure 3 panel A-B with Figure 4 panel B).

**Figure 4.**
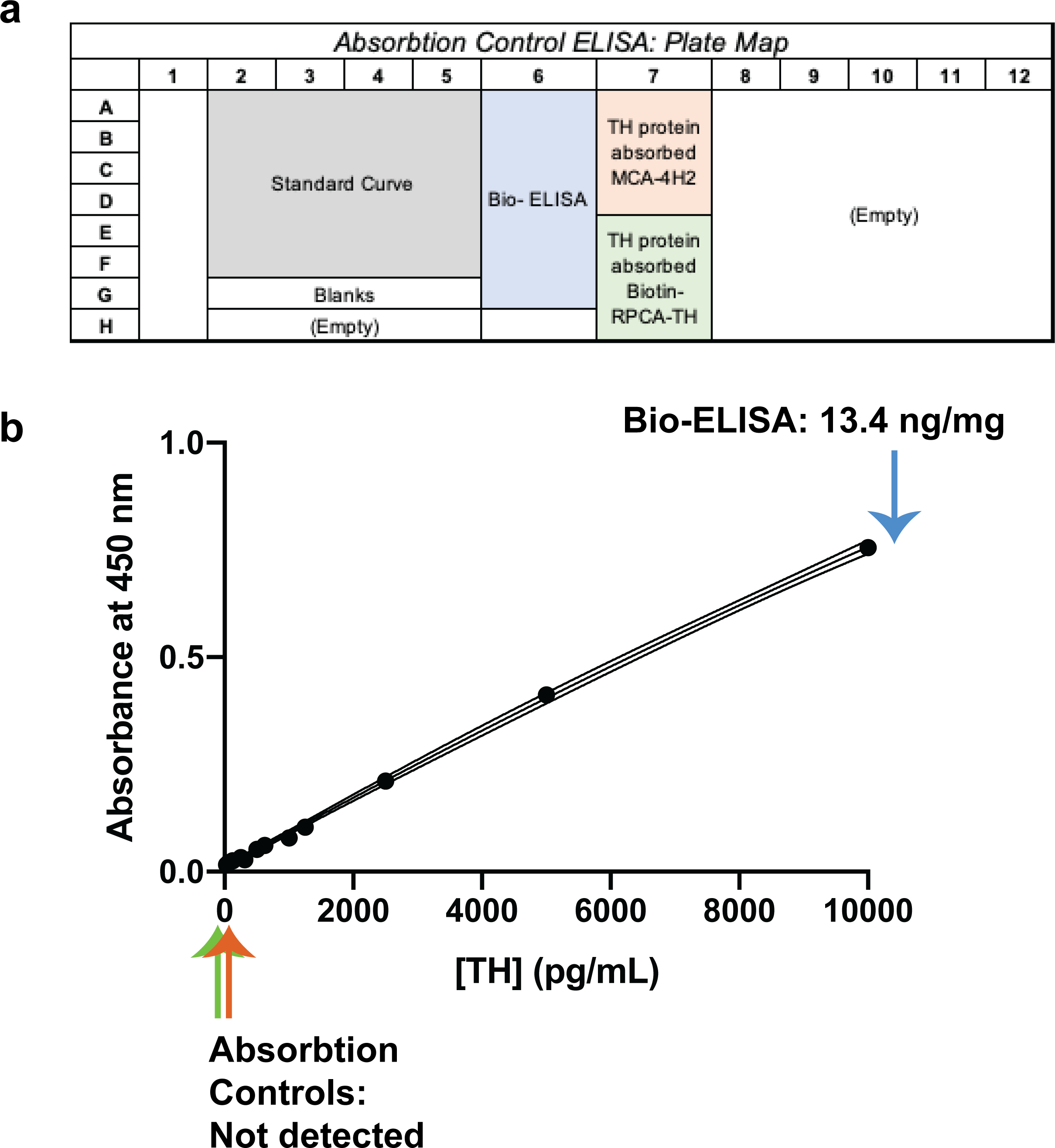
Absorption controls demonstrate specificity of TH Bio-ELISA. A) Schematic layout of experimental conditions to assess absorption controls in contrast to optimized Bio-ELISA conditions using PC12 cell lysate. B) Representative standard curve shown to illustrate PC12 cells’ TH concentration using optimized Bio-ELISA (blue arrow), absorbed capture antibody (MCA-4H2 preincubated with 20ug/mL recombinant TH, orange arrow), and absorbed detection antibody (biotinylated RPCA-TH preincubated with 20ug/mL recombinant TH, green arrow). PC12 TH is undetectable after absorption of either capture or detection antibodies, confirming assay specificity.

#### Contrary to our hypothesis, monocytes isolated from blood of PD patients show increased TH protein relative to age-matched healthy controls

PD is a disease in which monoamine signaling is affected in both CNS and peripheral immune cells^9^. The literature supports the hypothesis that similar to the CNS, peripheral TH expression is altered, but there is no reliable information about the direction of this change. Since peripheral immune cells including PBMCs express the machinery for catecholamine synthesis, including TH, they provide a biologically relevant peripheral tissue preparation to investigate TH levels in monocytes of PD patients and age-matched healthy subjects. Monocytes for each subject were isolated from 20 million total peripheral blood mononuclear cells (PBMCs) using anti-CD14 magnetic isolation per manufacturer’s instructions. Purified monocytes were immediately lysed and assayed *via* Bio-ELISA for TH concentration following total protein quantification. Of 11 healthy control samples included, only three registered TH concentrations above the detection threshold. By contrast, all 11 PD patients recruited for this study show clear positive TH values that were significantly higher than healthy controls. These data suggest that, contrary to our original hypothesis, PD monocytes express significantly more TH protein relative to healthy control subjects (Fig 5A - n=11, t[20]=3.777, P=0.0012). Mean TH protein concentrations in PD monocytes are shown on a representative standard curve (Figure 5B), along with raw data used to calculate TH concentrations (Figure 5C). While these data represent a snapshot of TH levels in circulation PD monocytes, we cannot make any overarching claims that TH levels in monocytes precede clinical symptoms of PD, or predict a PD diagnosis. A larger sample numbers and longitudinal studies can test these possibilities. Nevertheless, these data suggest that in peripheral monocytes of Parkinson’s patients, the rate limiting protein involved in catecholamines synthesis is increased. Investigating the potential mechanism was the focus of the next set of experiments.

**Figure 5.**
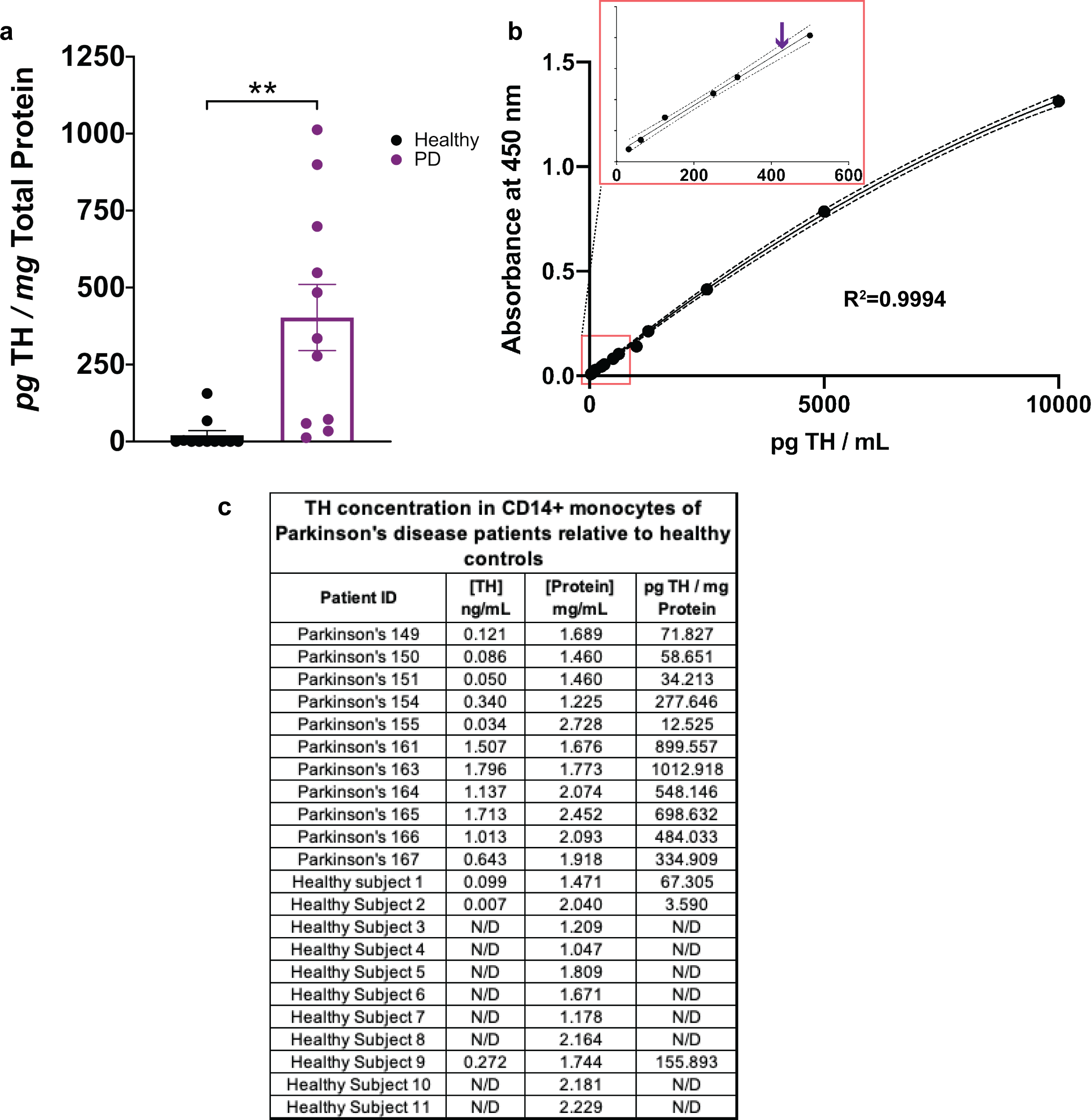
TH protein is increased in CD14+ monocytes isolated from PD patients. Total CD14+ monocytes were magnetically isolated from 20 million freshly isolated PBMCs derived from whole blood of 11 PD patients and 11 healthy volunteers, immediately lysed in the presence of protease inhibitor and stored at -80°C. Following protein quantification, whole lysate from each sample was added to duplicate wells and assayed for concentration of TH. A) Monocytes isolated from PD patients express significantly greater quantity of TH compared to equivalent monocytes isolated from healthy control subjects (unpaired two-tailed T-test, alpha=0.05, p<0.05) B) Mean TH concentration for monocytes from PD patients plotted on a representative standard curve, with inset magnifying the lower end of the curve. C) Calculations are shown by which raw TH concentration in ng/mL is normalized to total protein per sample. Samples included in A, each an independent biological replicate, are shown in C. Data are shown as ±SEM.

#### TNF**δ** increases number of TH+ monocytes, and the amount of TH protein per monocyte

There is strong evidence in the literature for increased TNFδ in PD^27, 34–36^ including in the brain, cerebrospinal fluid, and serum of Parkinson’s patients^27^ as well as in Parkinsonian mice^37, 38^. These reports suggest that TNFδ plays a role in the often hypothesized peripheral inflammation in PD^74–79^, which is also documented in other inflammatory states including rheumatoid arthritis^80, 81^ and multiple sclerosis^7^, where TH expression is linked to TNFδ expression^7, 80, 81^.

Therefore, we tested the hypothesis that *ex vivo* stimulation of monocytes from healthy subjects with TNFδ stimulates TH expression, as measured by changes in the number of TH-expressing monocytes, and/or the amount of TH per monocyte. We employed flow cytometry to address the former, and bio-ELISA to address the latter. Two million monocytes isolated from whole blood of healthy donors were treated for 4 hours with tissue plasminogen activator (TPA, 100ng/mL, positive control for increased monocyte TH expression^7^), TNFδ (17ng/mL)^61^ and compared with monocytes treated with vehicle (Figure 6A). Monocytes were assayed for TH expression by two complementary methods: flow cytometry^51^ (Figure 6B-F) and ELISA (Figure 6G-H). We should note that because a prolonged TNFδ exposure can induce cell toxicity^82–86^, we tested multiple treatment durations. We found that a 4-hour TNFδ (17ng/mL)^61^ treatment had a minimal effect on cell viability; whereas, a longer TNFδ exposure substantially decreased cell viability. Therefore, a 4-hour treatment strategy was selected in this study.

**Figure 6.**
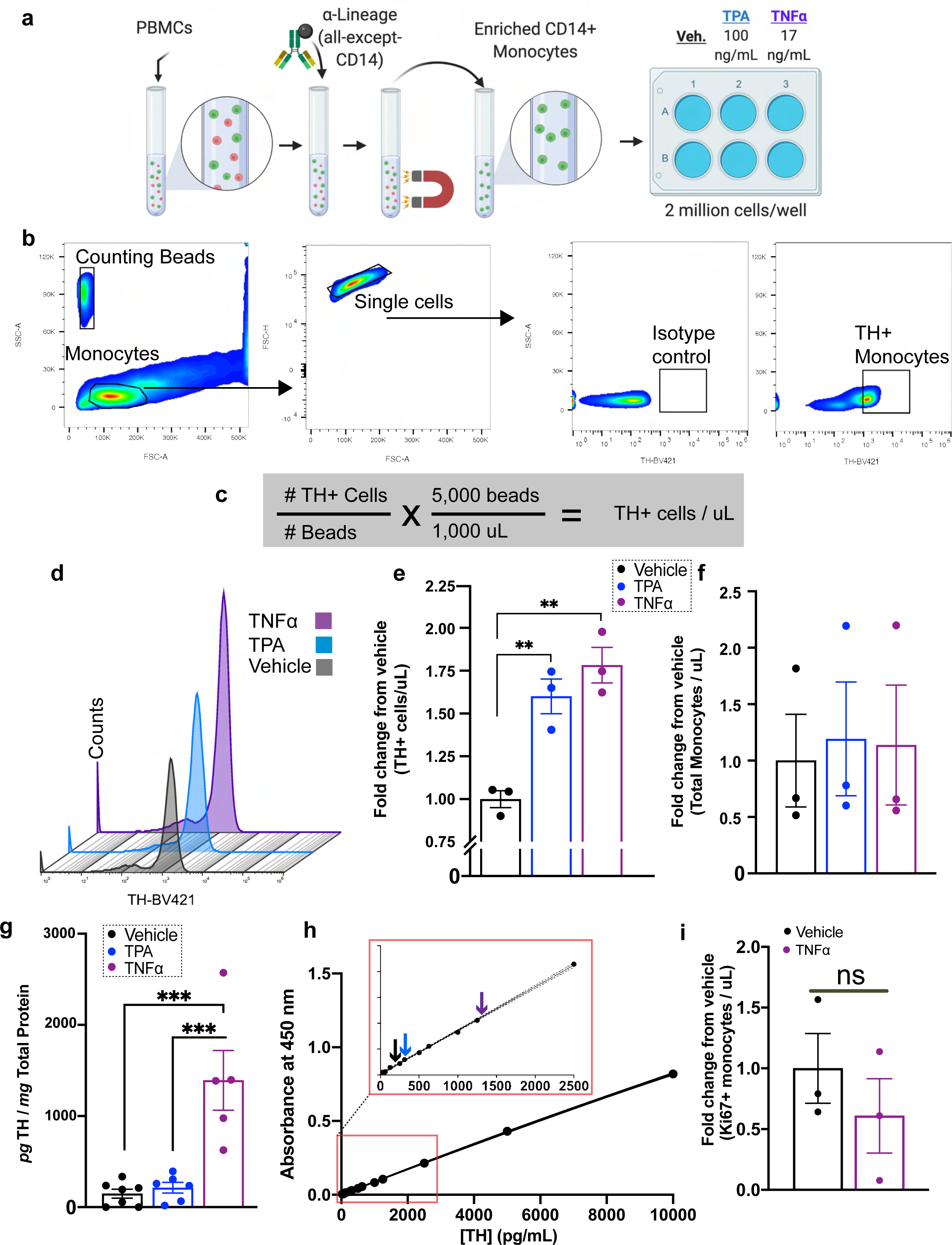
TNFδ increases number of TH+ monocytes and amount of TH protein per monocyte. A) Total CD14+ monocytes were isolated using negative magnetic selection from 80 million healthy donor PBMCs. Monocytes were seeded into a 6-well ultra-low-adherence plate at 2 million cells per well, and treated with vehicle (media), TPA (100ng/mL, positive control), TNFδ (17ng/mL), in duplicate. B) One duplicate was assayed by flow cytometry to detect TH-expressing monocytes, using counting beads as a reference value to quantify the number of TH+ cells. C) Number of TH+ cells were quantified as shown. D) Representative histogram showing one set of samples assayed for TH expressing monocytes following stimulation. E) Both TPA and TNFδ induced significant increases in TH-expressing monocytes relative to vehicle, shown as fold-increase relative to vehicle (n=3 per group, one-way ANOVA, p<0.01). F) No increase in total monocytes per condition, relative to vehicle (n=3 per group, one-way ANOVA, n.s.). G) TH concentration in picograms per milligram total protein shows TNFΑ treatment results in significantly increased TH protein relative to vehicle and TPA (n=5-6 per group, one-way ANOVA, p<0.001). H) Mean TH protein level for monocytes treated with vehicle, TPA and TNFδ are plotted on a representative standard curve, with the inset magnifying the lower end of the curve. I) Intracellular flow cytometry for Ki67 does not reveal significant differences between vehicle and TNFδ treatment groups, confirming a lack of cell proliferation following TNFδ treatment. Data are shown as ±SEM.

To control for donor variability, we added identical quantities of counting beads as a reference. The number of TH+ monocytes was quantified by flow cytometry (Figure 6B, left) in two experimental groups: TPA-treated and TNFδ-treated. Monocytes in each condition were gated to isolate single cells expressing TH (Figure 6B). Raw counts of monocytes in each condition revealed increased monocytes expressing TH after treatment with TPA or TNFδ (Figure 6D), while the number of TH+ monocytes per microliter (Figure 6C) are significantly increased relative to vehicle (Figure 6E; N=3, F(2,6)=0.364, p=0.0018), suggesting that the number of TH+ monocytes increase following treatment with TNFδ or positive TPA control. A possible mechanism for this observation is either monocyte proliferation during the treatment period or an altered monocyte phenotype in response to TNFδ, with no change in total number of monocytes. In other words, following TNFδ treatment, TH+ monocytes may either be increasing in number (proliferation) or existing monocytes upregulate TH expression and become TH+ (phenotypic change). While a four-hour exposure to TNFδ is an insufficient time period to induce proliferative events in immune cells^74^, we could not confidently rule out these possibilities without additional analyses. Therefore, we quantified monocyte proliferation by comparing the total number of monocytes per microliter of untreated vs. TNFδ treated experimental group. We found the total number of monocytes per microliter to be unchanged (Figure 6F). While these results suggest that monocyte proliferation did not occur in response to TNFδ, the results of simple cell counts are not definitive. We elected to take a more rigorous approach and assess Ki67 expression as a measure of cell proliferation^87^. Ki67 expression in TNFδ treated monocytes relative to vehicle-treated controls revealed no change in Ki67 expression following TNFδ treatment (Figure 6I). Thus, these results showed phenotypic changes in monocytes, but not cell proliferation in response to TNFδ. While this finding explains our earlier observation of increased numbers of TH+ cells, the potential phenotypic shift following TNFδ mediated immune stimulation was an unpredicted and novel finding.

Our flow cytometry data strongly support the conclusion that TNFδ increases numbers of TH+ monocytes, but increased number of TH+ monocytes could be due to increased numbers of cells expressing TH protein, increased quantity of TH protein per cell, or both. In order to determine whether or not TNFδ treatment increases quantity of TH protein per monocyte, identically treated monocytes were lysed and assayed using our TH Bio-ELISA. We found that four-hour treatments with TNFδ significantly increased the amount of TH protein (picogram TH per milligram total protein) above both vehicle and the positive control group (TPA treatment; Figure 6G; n=5-6 per group, F(2,15)=3.297, p=0.0001), indicating that exposure to TNFδ is sufficient to increase TH protein in human monocytes. Overall, our data show that TNFδ increases both the number of monocytes expressing TH and the quantity of TH expressed by each cell.

#### Inhibition of TNFδ blocks increase in number of TH+ monocytes and amount of TH per monocyte

To determine the specificity of TNFδ regulation of TH in monocytes, we employed two approaches. We investigated whether inhibition of TNFδ signaling attenuates or blocks the TNFδ mediated increase in TH. In addition, we asked whether or not interleukin-6 (IL6), a cytokine with pleiotropic effects ^71^ that is also increased in PD^88–90^, and is associated with non-motor symptoms of PD ^88–90^ can also regulate TH expression in the peripheral monocytes. To test these possibilities, we investigated whether XPro1595, a TNFδ inhibitor^91, 92^, reduces monocyte TH expression relative to TNFδ treatment alone. In parallel experiments, monocytes were treated with IL6. Two million monocytes isolated from whole blood of healthy donors (Figure 7A) were treated with XPro1595 alone (50ng/mL), TNFδ (17ng/mL) or TNFδ plus XPro1595 (Figure 7B), IL6 alone (17ng/mL) or IL6 plus XPro1595. The cells were subjected to flow cytometry or Bio-ELISA. Consistent with the literature ^71^, we found relative to TNFδ treatment alone, XPro1595 inhibition of TNFδ reduced both the number of TH+ monocytes and the quantity of TH per monocyte (Figure 7C and D), suggesting that soluble TNFδ mediates increased TH in human monocytes. As shown in Figure 7C and 7D, IL6 neither changed the number of TH+ monocytes nor the quantity of TH per monocyte. We should note that our data show that TNFδ is capable of regulating TH in monocytes whereas other elevated cytokines, including IL6, are not. Since we have not tested the effect of additional, non-upregulated cytokines, we cannot claim that what we have shown in this study is exclusively mediated by TNFδ. Instead, we only claim that TNFδ is capable of regulating monocytic TH. In addition, while have not investigated the direct link between increased TH protein in PD monocytes and TNFδ, our *ex vivo* data (Figure 6) support the interpretation that TNFδ plays a role in increased TH expression in immune cells of PD patients.

**Figure 7.**
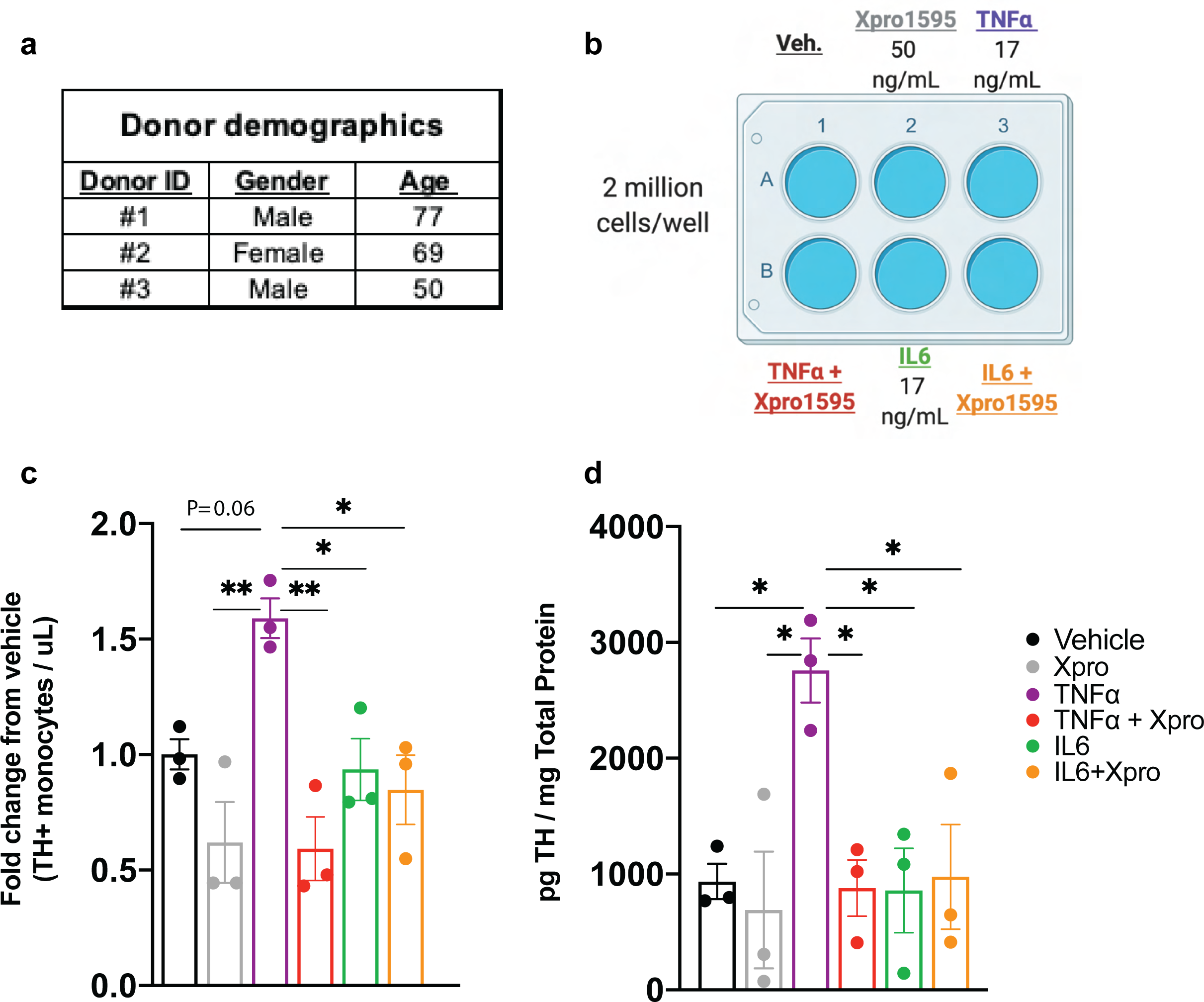
Inhibition of TNFδ blocks increase in number of TH+ monocytes and amount of TH per monocyte. A-B) Acutely isolated monocytes from three healthy donors were seeded at 2 million cells per well in duplicate ultra-low-adherence plates, and treated with TNFδ (17ng/mL), XPro1595 (50ng/mL), IL6 (17ng/mL) or combinations thereof as indicated. C) In samples assayed by flow cytometry, using counting beads as a reference value to quantify the number of TH+ cells, co-incubation with TNFδ and XPro1595 significantly reduced the number of TH+ monocytes relative to TNFδ treatment alone. Treatment with IL6 or IL6 + XPro1595 resulted in no significant change in the number of TH+ monocytes. Values are represented as fold change relative to vehicle (n=3 per group, one way ANOVA, *P<0.05, **P<0.01). D) TH concentration (picogram per milligram total protein) significantly increases upon TNFδ treatment, and is reduced significantly to baseline levels following co-incubation with TNFδ and XPro1595. Neither IL6 nor IL6+XPro1595 significantly increased TH quantity (n=3 per group, one way ANOVA, *P<0.05). Data are shown as ±SEM.

In summary, we developed a highly reproducible and quantitative Bio-ELISA to measure TH protein levels in murine and human cells. Following validation of our assay in multiple TH expression systems, we investigated TH expression in PD immune cells and of age-matched healthy control subjects. We observed that PD patients’ monocytes expressed significantly greater amounts of TH per monocyte. Inspired by the literature indicating increased TNFδ in PD, we uncovered an intriguing link between TNFδ stimulation and increased TH expression in healthy monocytes, which is attenuated by treatment with TNFδ inhibitor Xpro1595. Given that TH expression and catecholamine release has been shown to be associated with an anti-inflammatory effect and can mitigate TNFδ mediated inflammation, we posit that increased TH expression in monocytes in response to elevated TNFδ is a compensatory mechanism. This observation is a step towards understanding the potential underlying mechanism and functional consequence of changes in catecholamines in peripheral immune system in PD. Nevertheless, we acknowledge that one of the limitations of this study was the infeasibility of quantifying TNFδ responses in PD monocytes. Whereas, TH can be quantified in a 30 mL blood sample (Figures 3-5), for functional assays (Figures 6 and 7) a large blood volume (∼500 mL) is required, which is not feasible in PD subjects. In addition, our data represent merely a snapshot of TH levels present in circulating PD monocytes at a single timepoint; we do not make any claims that elevated levels of TH expressing monocytes precede or predict PD. Larger sample numbers and longitudinal studies can test these possibilities. Nonetheless, the current results raise many interesting questions: Do the circulating TH expressing monocytes reflect changes in central dopamine? Does effective PD therapy reduce the level of TH in peripheral immune cells? Does elevated TH in monocytes predict PD onset or its progression? Future studies will examine these questions and the connections between the peripheral immune system to the brain.

## Methods

### Human subjects

Human brain tissues were obtained *via* approved IRB protocols #IRB201800374 and IRB202002059 respectively. Blood samples were obtained at the University of Florida Center for Movements Disorders and Neurorestoration according to an IRB-approved protocol (#IRB201701195).

### Brain tissues from healthy subjects

Human brain tissues were obtained *via* approved IRB protocols IRB202002059 and IRB201800374, from the UF Neuromedicine Human Brain and Tissue Bank (UF HBTB). The tissues were not associated with identifying information, exempt from consent, therefore no consent was required. Regions of interest were identified and isolated by a board-certified neuropathologist.

### Blood samples from healthy subjects

Blood samples from age-matched healthy subjects were obtained from two sources: an approved IRB protocol with written informed consent (IRB201701195), or were purchased from Lifesouth Community Blood Center, Gainesville, FL from August 2017 to January 2020 as deidentified samples, and exempt from informed consent (IRB201700339). According to Lifesouth regulations, healthy donors were individuals aged 50-80 years-old of any gender, who were not known to have any blood borne pathogens (both self-reported and independently verified), and were never diagnosed with a blood disease, such as leukemia or bleeding disorders. In addition, none of the donors were using blood thinners or antibiotics, or were exhibiting signs/symptoms of infectious disease, or had a positive test for viral infection in the previous 21 days.

### Blood samples from PD patients

Blood samples were obtained from PD patients (aged 50-80 years-old of any gender) at the University of Florida Center for Movements Disorders and Neurorestoration according to an IRB-approved protocol (#IRB201701195), via written informed consent. All recruited patients’ PD was idiopathic. Patients did not have any recorded blood-borne pathogens or blood diseases, nor were they currently taking medications for infections according to their medical record. In addition, none of the donors were using blood thinners (warfarin, heparin), antibiotics, over-the-counter (OTC) medications other than aspirin, or were exhibiting signs/symptoms of infectious disease or had a positive test for viral infection in the previous 21 days. Current medications are summarized in Supplementary Table 1.

### TH recombinant protein

Full length human TH protein was expressed from a synthetic cDNA inserted into the *EcoRI* and *SalI* sites of the pET30a(+) vector and was codon optimized for expression in *E. coli*. The vector adds an N-terminal His-tag and other vector sequence, a total of 5.7kDa. Expression of the construct was made by standard methods and purification was performed using the His tag by immobilized metal affinity chromatography on a nickel column. The TH sequence used in this study is the human tyrosine 3-monooxygenase isoform shown in Uniprot entry P07101-2.

### Model systems used for the validation of bio-ELISA

#### Human macrophages

Primary human macrophages were cultured as described previously^43^. Peripheral blood mononuclear cells (PBMCs) isolated as described below were re-suspended in RPMI 1640 containing 1% Pen/Strep and 7.5% sterile-filtered, heat-inactivated autologous serum isolated from the donor’s own blood, and plated in 24-well untreated polystyrene plates at 1 million PBMCs per well. To retain only monocytes/macrophages, cells were washed after 90 minutes of adherence time to remove non-adherent cells with incomplete RPMI 1640, followed by replacement with complete media. Media was replaced at days 3 and 6 following culture, and cell lysis performed on day 7 following culture.

#### Primary murine midbrain dopamine neurons

Midbrain dopamine neurons strongly express TH^44^ and were used as a positive control group. Acutely dissociated mouse midbrains from 0-2 day-old male and female pups were isolated and incubated in dissociation medium at 37°C under continuous oxygenation for 90 minutes. Dissociated cells were pelleted by centrifugation at 1,500×*g* for 5 min and resuspended and triturated in glial medium (Table 1). Cells were then plated on 12 mm coverslips coated with 0.1 mg/ml poly-D-lysine and 5 μg/ml laminin and maintained in neuronal media. Every 4 days, half the media was replaced with fresh media. The materials used for the preparation and maintenance of midbrain neuronal culture are outlined in Table 1.

#### Positive and negative control cell lines

All cell cultures were maintained at 37°C with 5% CO2 and all cell culture supplies are listed in Table 2. HEK293 cells^45^ are not thought to express TH and so were used as a negative expression control and were cultured as described previously^46, 47^. PC12 cells express TH^48^ and were used as a positive control. The cells were cultured as described by Cartier et al. 2010^49^. CHO cells were cultured as previously described^50^, and were used as a negative control for TH expression.

### PBMC isolation

PBMCs express TH^43, 51^. As previously published^51^, whole blood was collected in K2EDTA vacutainer blood collection tubes (BD, 366643) and held at room temperature for up to 2 hours prior to PBMC isolation. Briefly, blood from healthy volunteers and PD patients was overlaid in Leucosep tubes (Table 2) for PBMC isolation, centrifuged for 20 minutes at 400g with brakes turned off and acceleration set to minimum. PBMCs were collected from the interphase of Ficoll and PBS, transferred to a fresh 15mL conical tube, resuspended in 8mL sterile PBS and centrifuged for 10 minutes at 100g, and repeated twice more. Cells were counted with a hemacytometer using trypan blue exclusion of dead cells, and density-adjusted for downstream applications.

### Magnetic monocyte isolation

PBMCs are composed of multiple cell subsets^52^, each with distinct function and catecholamine sensitivity^53, 54^ – for example, lymphocyte regulation by catecholamines dopamine and NOR^5, 6, 55^ have been studied for several decades^8, 18, 56, 57^, while data regarding catecholamine function in myeloid lineage cells including monocytes is less abundant. In this study, we were narrowly focused on studying peripheral monocytes which we and others have previously shown to express TH^9, 51, 58–60^. Because PBMCs comprise a variety of immune cell types, we used immunomagnetic enrichment to obtain a greater than 95% CD14+ monocytes that were utilized in assays described in the current study. Supplementary Figure 2 shows representative flow cytometry data from routine verification of monocyte enrichment. (Supplementary Figure 2).

CD14+ monocytes express TH^51^. Primary CD14+ monocytes were isolated using Biolegend MojoSort magnetic isolation kit (Biolegend, 480094) per manufacturer’s instructions. Briefly, 20 million total PBMCs were counted, density adjusted to 1 million cells/uL, resuspended in MojoSort buffer, and incubated with TruStain Fc-block for 10 minutes at room temperature, followed by 1:10 anti-CD14 magnetic nanobeads for 15 minutes on ice. Following 2 washes with 2.5mL ice-cold MojoSort buffer, cell pellet was resuspended in 2.5mL MojoSort buffer and subject to three rounds of magnetic isolation per manufacturer’s instructions. The resulting cell pellet was washed to remove remaining non-CD14+ cells and subject to cell lysis as detailed below.

### Preparation of cell lysates

Adherent cells in culture were lifted using 0.02% EDTA in PBS, diluted with 5 volumes of PBS, and centrifuged at 100 x *g*. Non-adherent cells (PC12) were centrifuged at 100 x *g* for 5 minutes at room temperature, and cell pellets were washed 3 times with 5 volumes of sterile PBS. Primary macrophages and primary murine neuron cultures were washed thrice with ice-cold PBS, on ice. Cell pellets and adherent primary cells were then lysed in ice-cold lysis buffer (10mM NaCl, 10% glycerol (v/v), 1mM EDTA, 1mM EGTA, and HEPES 20mM, pH 7.6), with Triton X-100 added to a final concentration of 1%, containing 1x protease inhibitor cocktail (Millipore-Sigma, 539131) for one hour at 4°C with rotation. Resulting lysate was centrifuged at 12,000 x *g* for 15 minutes at 4°C. Supernatant was set aside for protein quantification by Lowry assay (Biorad, 5000112) and the remainder was stored at -80°C until use for downstream assays.

### Western blot

Reagents, antibodies and equipment are outlined in Tables 2, 3 and 4. Samples of PC12 lysate (5ug) and recombinant TH protein (120ng, 60ng, 30ng, 15ng, 7.5ng, 3.75ng, and 1.875ng) were incubated in Laemmli sample buffer containing 10% beta-mercaptoethanol at 37°C for 30 minutes, separated by SDS-PAGE on 10% bis/polyacrylamide gels, and transferred to nitrocellulose membranes. After first blocking for 1 hour in TBS-T (50mM Tris-HCl, 150mM NaCl, and 0.1% Tween 20) containing 5% dry milk (blocking buffer), then incubated with primary antibody against TH (Table 4) overnight at 4°C. Membranes were then incubated with an appropriate secondary antibody (Table 4) for 1 hour at room temperature with agitation.

Following all antibody steps, membranes were washed three times for 5 minutes each using TBS-T. TH was visualized using the Licor Odyssey (Table 2). Absorption controls were performed as followed: the primary antibodies were pre-incubated with 20ug/mL recombinant TH protein for 30 minutes on ice, then were used to confirm primary antibody specificity (Table 3, Figure 2C-D).

### Immunohistochemistry

Human tissues were sectioned at 40µm on a vibrating microtome and subjected to antigen retrieval in citrate buffer (10mM citric acid, 2mM EDTA, 2% Tween-20, pH 6.2) at 96°C for 30 minutes, and then allowed to cool to room temperature. PFA-perfused mouse brain tissues were also sectioned at 40 µm on a vibrating microtome.

Human and murine brain tissues were quenched for 20 minutes with 3% hydrogen peroxide, blocked and permeabilized at 37°C for 1 hour in PBS containing 5% normal goat serum and 0.5% TritonX-100. Primary antibodies RPCA-TH and MCA-4H2 (1:500 and 1:100 dilution, respectively, Table 4) were incubated overnight, followed by secondaries conjugated to HRP (1:250, Table 4), incubated for 1 hour at room temperature. Isotype control antibodies (Biolegend, Table 1) were used to confirm specificity of RPCA-TH and MCA-4H2. Sections were detected with HRP-substrate NiDAB (Vector Labs, Table 3).

### Detection antibody (RPCA-TH) Biotinylation

EZ-Link Sulfo-NHS-LC-Biotin (A39257, Thermo Scientific) at 20-fold molar biotin was used according to the manufacturer’s protocol. Anti-biotin antibody was concentrated to 2mg/mL, pH was adjusted to 8.0 at room temperature. The conjugate was purified by gel filtration on a Biorad 10DG column (cat 732-2010) at room temperature.

### ELISA for TH

Antibodies used for ELISA are described in Table 1. Ten lanes of an Immulon 4 HBX High-Binding 96 well plate were coated with 100uL per well of 1:1,000 dilution of 1mg/mL mouse anti-TH (MCA-4H2) in coating buffer (28.3mM Na_2_CO_3_, 71.42mM NaHCO_3_, pH 9.6) for 20 hours at 4°C. Edge lanes 1 and 12 were left empty. Wells were blocked with 5% fat free milk in 1x TBS (pH 7.4) for 1 hour at room temperature on an orbital shaker set to 90rpm. To produce a standard curve, two standard curve lanes were generated, with six serial dilutions, beginning at 10ng/mL and 1ng/mL in TBS-T containing 1% fat free milk (with the last well in each standard curve lane left with incubation buffer only as a blank. Remaining wells were incubated in duplicate with 100 microliters of lysates from 1.5 million cells of interest. Incubation was completed for 20 hours at 4°C on an ELISA shaker set to 475 rpm.

After each well was washed and aspirated 6 times with TBS-T, affinity purified polyclonal rabbit anti-TH (EnCor, RPCA-TH) conjugated to biotin was diluted 1:6,000 from a stock concentration of 1.65mg/mL in TBS-T with 1% fat-free milk and incubated for 1 hour at room temperature at 425rpm. 100uL Avidin-HRP (Vector labs, A-2004), diluted 1:2,500 in TBS-T with 1% fat-free milk, was added to each well following washing as described above, and incubated for 1 hour at room temperature at 425rpm. Following final washes, 150uL room temperature TMB-ELISA reagent (Thermo Fisher, 34028) was added to each well. The reaction was allowed to continue for 20 minutes, protected from light, and stopped by addition of 50uL 2N H_2_SO_4_. The plate was immediately read at 450nm. Absorption controls (Figure 4) were conducted by pre-incubating MCA-4H2 and RPCA-TH with a 20-fold excess concentration of recombinant TH protein for 30 minutes on ice, prior to addition to the ELISA plate, followed by the remainder of the protocol described above.

Duplicate standard and sample wells were averaged, and background-subtracted based on blank wells. The concentration of TH for each experimental group was calculated using a quadratic curve equation calculated in Graphpad Prism 8, then normalized to total protein concentration per sample as calculated using the Lowry assay. Samples which produced negative values for TH concentration were considered below detection threshold, and therefore assigned a value of 0. Final TH values shown are presented as pg TH/mg total protein after multiplication of the nanogram TH value by 1,000 to show TH as picogram TH/milligram total protein.

### *In vitro* stimulation/treatment with TNF**δ**, tissue plasminogen activator (TPA), TNF**δ** inhibitor XPro1595 and IL6

Monocytes were isolated from total PBMCs prepared as described above^51^ using negative selection (Biolegend, 480048) per manufacturer’s instructions. Total PBMCs were Fc-blocked to reduce nonspecific binding, followed by incubations with biotin-conjugated antibody cocktail containing antibodies against all subsets except CD14 (negative selection), followed by incubation with magnetic-Avidin beads, allowing all subsets other than CD14+ monocytes to be bound to the magnet. Monocyte purity/enrichment was routinely verified to confirm that the final cell population was greater than 95% pure CD14+ cells (Figure S2). CD14+ monocytes were collected from the supernatant fraction, washed, counted and density adjusted such that 2 million CD14+ monocytes were seeded per well (Figure 5A) and treated for 4 hours with vehicle, TPA (100ng/mL, Biolegend, 755802)^7^, TNFδ (17ng/mL, Biolegend, 570102)^61^, XPro1595 (50ng/mL), or IL6 (17ng/mL) in an ultra-low-adherence 6-well plate (Corning, 3471) to prevent adherence. Suspended cells from each treatment group were aspirated and placed in a 15mL conical tube, with any remaining adherent cells detached by incubation with 700uL Accumax solution for 3 minutes (Innovative Cell Technologies, AM105) and added to suspended cells. After pelleting cells by centrifugation (3 minutes x 100g, room temperature), cells were assayed by either flow cytometry^51^ or lysed for ELISA as stated above (“Preparation of cell lysates”).

As previously published^51^, cells for flow cytometry were fixed and permeabilized (eBioscience, 88-8824-00), and stained for intracellular marker TH (Millipore-Sigma, AB152, 1:100) followed by a species-specific secondary (anti-Rabbit BV421, BD, 565014). After resuspending the sample in a final volume of 250uL PBS, 5uL of Invitrogen CountBright Absolute Counting Beads (5000 beads/mL, Invitrogen, C36950) were added just prior to data acquisition (Sony Spectral Analyzer, SP6800). Monocytes were gated for single cells and positive TH expression (Figure 5B), and normalized to counting beads in each sample to obtain an absolute count of TH+ monocytes per uL suspension.

### Statistics

A two-tailed, unpaired T test was used to compare TH quantity in PD patients versus healthy control. In this experiment, P<0.05 was considered statistically significant. One-way ANOVA with Tukey’s correction for multiple comparisons was used to compare TH-expressing monocytes assayed by flow cytometry and ELISA following treatment with TPA, TNFδ, XPro1595, IL6 or Vehicle. P<0.05 was considered statistically significant.

## Data Availability

All data will be made available upon reasonable request.

## Acknowledgements

This work was funded by T32-NS082128 (to A.G.), National Center for Advancing Translational Sciences of the National Institutes of Health under University of Florida Clinical and Translational Science Awards TL1TR001428 and UL1TR001427 (to A.G.), R01NS071122-07A1 (to H. K.), NIDA Grant R01DA026947-10, National Institutes of Health Office of the Director Grant 1S10OD020026-01 (to H. K.), UF-Fixel Institute Developmental Fund, DA043895 (to H.K.), by the University of Florida McKnight Brain Institute (MBI) (to A.G.), by the Bryan Robinson Foundation (to A.G.) and by The Karen Toffler Charitable Trust (to A.G.).

## Author Contributions

M.F., M.B., G.S., A.C., D.R.M., C.A.H., A.D, and A.G., performed experiments, contributed written portions, figures and methods to this manuscript. P.M., M.G.T., I.M., A.R.Z., M.S.O., W.J.S., H.K., and A.G., supervised, designed experiments in addition to direct contributions to the manuscript.

---

**Supplementary Figure 1.**
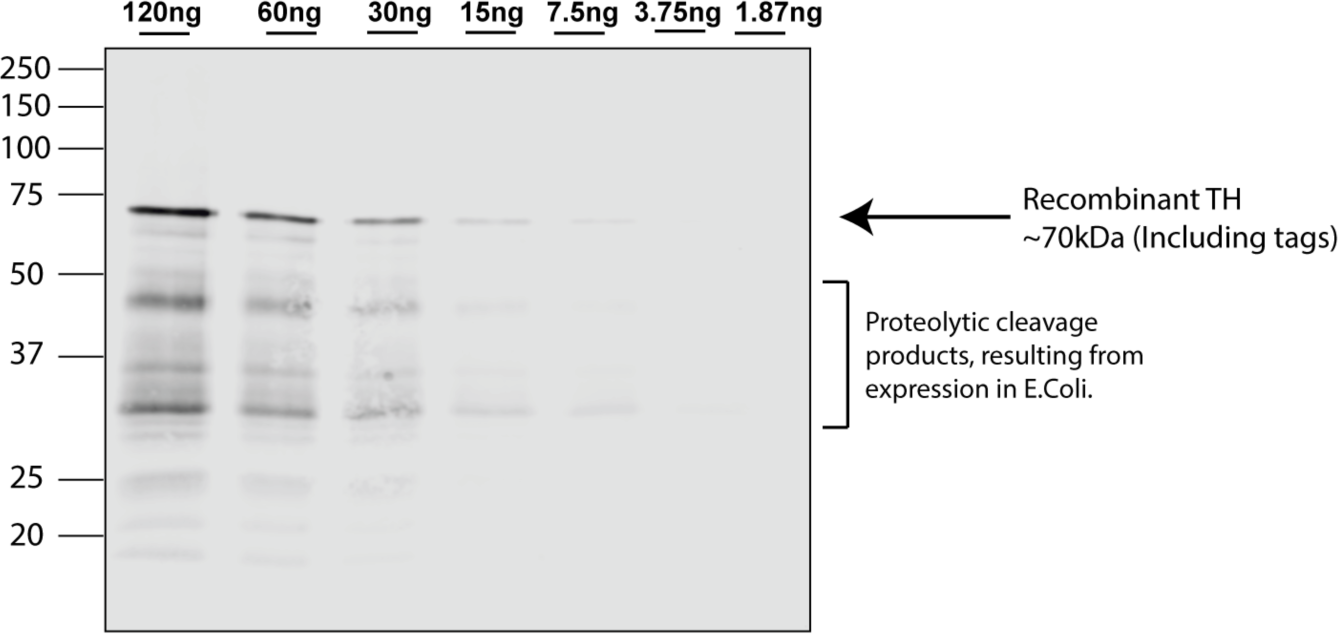
Lower molecular weight bands in recombinant TH protein are proteolytic cleavage products resulting from prokaryotic expression of TH protein. Similar to lower molecular weight bands seen in Figure 2, proteolytic cleavage is evident in recombinant TH protein assayed by western blot.

**Supplementary Figure 2.**
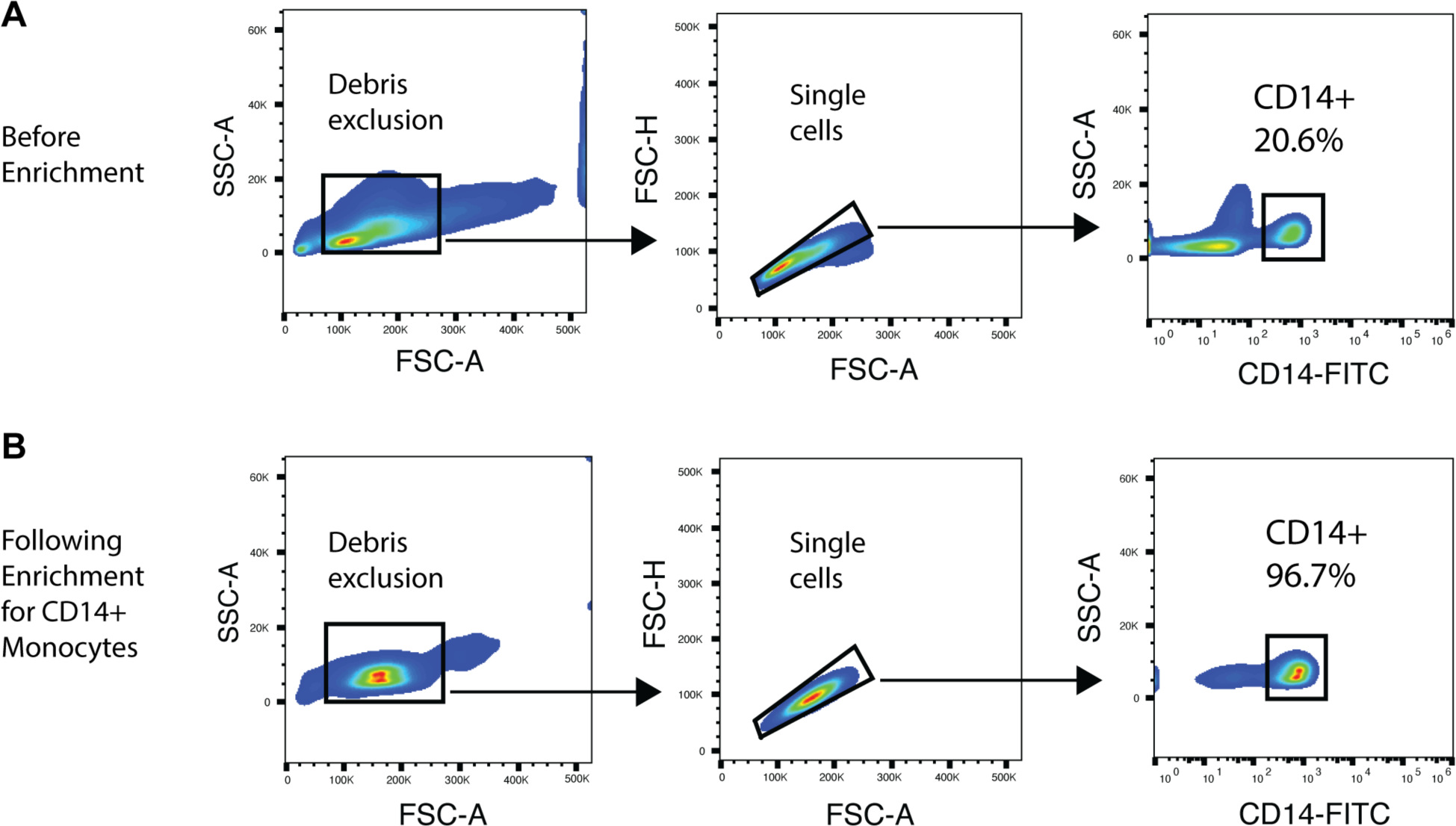
Enrichment for CD14+ monocytes from total PBMCs. A) Total PBMCs isolated from healthy donor whole blood shows ∼20% CD14+ monocytes as a fraction of total PBMCs. B) Following magnetic selection (see methods), a highly enriched population of monocytes consisting of greater than 95% CD14+ monocytes is available for downstream cell culture treatments.

**Supplementary Table 1:** Clinical data for Parkinson’s disease patients. All PD patients and healthy control subjects were free from blood borne pathogens, viral/bacterial infections, had not been treated for infections within the preceding 21 days and were not taking blood thinners other than aspirin. Detailed medical histories for healthy control subjects were not available, other than those data presented. UPDRS Off represents Part 3 motor scores when subjects were not currently administered dopamine replenishment therapy (L-DOPA/Sinemet). UPDRS On represents Part 3 motor scores 30 minutes following dopamine replenishment therapy (L-DOPA/ Sinemet).

| Supplementary Table 1: Clinical data for Parkinson's disease patients |  |  |  |  |  |  |  |  |
| --- | --- | --- | --- | --- | --- | --- | --- | --- |
| Patient ID | Disease Duration (Years) | Sex | Age (Years) | H-Y | UPDRS Off | UPDRS On | Other Conditions | Medications |
| Parkinson's 149 | 7 | M | 71 | 2 | 33 | -- | None | Rytary, Ropinirole, Amantadine, Azilect, Apokyn |
| Parkinson's 150 | 9 | M | 50 | 2 | -- | 26 | None | Sinemet, Selegiline, Mirapex |
| Parkinson's 151 | 19 | F | 70 | 2 | 38 | 37 | RLS | Sinemet, Pramipexole |
| Parkinson's 154 | 15 | M | 76 | -- | -- | -- | None | Asprin, Sinemet, Klonopin, Rasagilline, Rigotine |
| Parkinson's 155 | 9 | M | 85 | 3 | -- | 25 | HTN, T2D | Atorvastatin, Sinemet, Zetia, Losrtan, Metformin, Metoprolol, Flomax |
| Parkinson's 161 | 2 | M | 61 | 2 | -- | 33 | HTN | Amantadine, Sinemet, Atorvastatin, Losartan, Metoprolol, Omeprazol |
| Parkinson's 163 | 7 | M | 70 | 3 | 15 | -- | None | Sinemet, Amantadine, Flomax, Ropinirole |
| Parkinson's 164 | 4 | F | 67 | -- | 17 | -- | None | Atorvastatin, Sinemet, Fluoxetine, Ganapentin, Losartan |
| Parkinson's 165 | 1 | M | 60 | 3 | -- | 23 | Arthritis | Sinemet, Amantadine, Celebrex, Sertraline, Trazadone |
| Parkinson's 166 | 3 | F | 67 | 3 | -- | -- | None | Sinemet, Tylenol |
| Parkinson's 167 | 6 | M | 59 | 2 | -- | 26 | T2D | Sinemet, Asprin, Amlodipine, Metformin, Ibuprofen |
| Abbreviations: RLS (restless leg syndrome), HTN (hypertension), T2D (type 2 diabetes), H-Y (Hoen Yar), UPDRS (universal Parkinson's disease rating scale) |  |  |  |  |  |  |  |  |

